# Survey of Anisoptera diversity in flood prone areas in Tirurangadi taluk of Kerala, India

**DOI:** 10.1101/2020.06.28.168336

**Authors:** Arshad Kalliyil, MK Jasna Sherin, K Ashiba Amal, Nimna Noushad, P Shibitha, CT Hency, P Fathimath Sahla, Bini Mathew

**Affiliations:** Department of Zoology, PSMO College Tirurangadi, Malappuram, Kerala, India, 676306

## Abstract

Biodiversity forms the foundation of the vast array of ecosystem that critically contribute to human well being. Biodiversity is important in human managed as well as natural ecosystems. The changing status of insects and other invertebrates is the key indicator of biodiversity and environment that shapes it. The order Odonata comes under the class insecta. Odonata are divided into two suborders; Zygoptera (damselflies) and Anisoptera (dragonflies). Natural calamities can alter the ecosystem and thereby biodiversity drastically. This study was conducted in Tirurangadi taluk of kerala, India which is a flood prone taluk due to the over flow of the Kadalundi river during monsoon season. We reported a total of 12 species belonging to 10 genera and 2 families during the entire study. Sub-order Anisoptera was represented by the families, Libellulidae and Aeshnidae, comprising 11 species of Libellulidae and one species of Aeshnidae. On a long term, we aim at studying the impacts on insect biodiversity caused by floods.

## Introduction

Biodiversity is the variability among living organisms from all sources, including terrestrial, marine and other aquatic ecosystems and ecological complexes of which they are part; this includes diversity within species, between species and of ecosystems. Biodiversity forms the foundation of the vast array of ecosystem services that critically contribute to human well being. Biodiversity is important in human managed as well as natural ecosystems. The changing status of insects and other invertebrates is a key indicator of biodiversity and the environment that shapes it. Estimates of the total number of species or those within specific orders are often highly variable. Globally, averages of these predictions estimate there are around 5.5 million insect species with around 1 million insect species currently found. Very occasionally the record also show mass extinctions of insects. There are around 5680 species of odonata known from all over the globe. Many species often have small distributions and are habitat specific, hence they are often successfully used as indicators for environmental health and conservation management. Approximately 503 species of odonates occur in India with approximately 186 species being endemic. Endemic odonate fauna of India is largely concentrated in two biodiversity hotspots of India, viz., Western Ghats and north east India.

Insects are tracheate arthropods in which the body is divided into head, thorax and abdomen. A single pair of antennae (homologous with the antennules of the crustacea) is present and the head also bears a pair of mandibles and two pairs of maxillae, the second pair fused medially to form the labium. The thorax carries three pairs of legs and usually one or two pairs of wings. The abdomen is devoid of ambulatory appendages, and the genital opening is situated near the posterior end of the body. Postembryonic development is rarely direct and a metamorphosis usually occurs.

The name Odonata, derived from the Greek “odonto”-, meaning tooth, refers to the strong teeth ‘. found on the mandibles of most adults. Odonata are divided into two suborders: Zygoptera (damselflies) and Anisoptera (dragonflies).In Zygoptera front and hind wings are similar in shape and in Anisoptera hind wings are broader near the base than the front wings. Both are one of the most common insects flying over forest, fields, meadows, ponds and rivers. About 6,000 extant species are distributed all over the world. India is highly diverse with more than 500 known species. Odonata are one of the ancient orders of insects. It first appeared during the Carboniferous era, about 250 million years ago along with mayflies (Ephemeroptera). Odonata of the Carboniferous era consists of giants; for example *Meganeuropsis americana* from that era had a wingspan of 71 cm, almost the span of pigeon. Dragonflies and mayflies are ancient groups of insects, which amongst others, were the first to develop wings and venture into air. Dragonflies mastered the art of flying and continue to be the masters aerobats.

The life history of odonates is closely linked with water bodies. They use a wide range of flowing and stagnant water bodies. Even though most species of odonates are highly specific to a habitat, some have adapted to urban areas and make use of man-made water bodies. Habitat specificity has an important bearing on the distribution and ecology of odonates.

Dragonflies have a bullet shaped body and that helps them survive by being able to change directions fast. An adult dragonfly has three distinct segments, the head, thorax and abdomen as in all insects. The head has very short and inconspicuous, three to seven segmented antennae and has modified in association with great development of eyes, which cover two-third portion of its surface. The characteristic feature of dragonfly is its flying adaptation. It has fast wing flapping and that allows them to fly faster (about 55 miles per hour).Another neat feature of dragonflies’ wings are that they can help keep dragonflies warm. So if it’s a cool or cloudy day, they will shake their wings to warm themselves up. Unlike other insects, they do not perform any courtship, they copulate while in flight. As they exhibit sexual dimorphism males are easier to identify than their female counterparts.

Ovipositor in dragonflies may be either entophytic or helophytic. In the latter case the eggs are rounded and are either dropped freely into the water or attached superficially to aquatic plants. this method is the rule among the Anisoptera, with the exceptions mentioned below. In *Sympetrum* and *Tetragoneuria* the eggs are laid in gelatinous strings attached to submerged twigs. Endophytic oviposition is characteristic of Zygoptera and the Anisopteran families Aeshnidae and Petaluridae. Dragon flies adopting this method have elongate eggs which they insert into slits cut by the ovipositor in the stem and leaves of plants or other objects, near or beneath the water. In some cases the female (alone, or accompanied by the male) descends below the water surface for the purposer. Before the nymph emerges from the egg it swallows amniotic fluid, the associated contractions of stomodaeal musculature being visible through the shell. Pressure of the head of the embryo against the chorion is the immediate cause of hatching, since it forces open the lid like anterior extremity of the egg. the newly hatched insect is known as pronymph: at this stage it exhibits a more or less embryonic appearance, the whole body and appendages being invested by a delicate cuticular sheath. The pronymph is of extremely brief duration, lasting only a few seconds in a *Anax*(Tillyard), for two or three minutes in *coenagrion (B*alfour-Browne,1909), but upto 30 minutes in *Sympetrum striolatum*. At this stage the pulsations of the stomodaeum increase in frequency and the pronymphal cuticle is ruptured. The insect which emerges is in its second instar and is now a free nymph fully equipped for its future life.The nymph of the Odonata are campodeiform and may be divided into two main types- the Anisopteran and the Zygopteran. In the former the body ends in three small processes, a median epiproct or appendix dorsalis and a pair of lateral para procts: when closed they form a pyramid which conceals the anus. Respiration takes place by means of concealed rectal tracheal gills. In the Zygopteran type the three terminal processes are greatly developed to form caudal gills, and rectal tracheal gills are wanting.

*Megalagrion oahuense* nymphs are terrestrial, living among moist debris on the floor of Hawaiian forest and some other species of *Megalagrion* spend much time crawling out of streams in a water film on rocks. A few other terrestrial or semi - terrestrial nymphs (e.g. wolfe, 1953; Willey, 1955)but otherwise immature stage of the Odonata are exclusively aquatic, living in various situations in fresh water. Many remain hidden in sand or mud and are homogenously coloured without any pattern. Those which live on tyhed river-bottom or among weed exhibit a cryptic pattern which tends to conceal them from enemies and prey. The number of instars that intervene between the egg and the imago varies in different species and also among individuals of the same species. It ranges between about ten and fifteen and the whole nymphal period may be passed through within a year as in most Zygoptera, or occupie two years as in *Aeshna*, or may even last from three to five years. In temperate species there may be a diapauses, such as occurs in the last instar nymph of Anax *imperator* and which ensures the synchronised emergence of adults in the following spring.

Most of the records of longevity in nature refer only to the reproductive period. During this, most dragonflies live up to 6 weeks. If maturation period is included, it may extend up to 8-10 weeks, respectively. It is known that aestivating spread wings (Lestidae) can live much longer as adults. Dragonflies encounter a large number of predators throughout their life. Fishes are important predators during the larval stage. Birds such as Hobby (*Falco subbuteo*), Bee-eaters (*Merops sp.*), Kingfishers, Herons and Terns have been observed to feed on odonates. Large dragonflies, robberflies (Asilidae) and spiders are important invertebrate predators. Parasitizing females climb or swim beneath the water to search for the eggs in the submerged plants. Many migrating species are intermediate hosts of avian trematode parasites like *Prosthogonimus*. During mass emergence of these with the female while laying egg. Usually during this period the female is very vulnerable to the attack by other males. Non-mated males attack the mated pair and try to hijack the female. In such cases the hovering male anchors the egg-laying female. This predation forms an important link in the transfer metacercariae and cysts of the parasite. Larval stages of water mite (*Hydrachnidia*) parasitise odonates.

Odonates, being predators both at larval and adult stages, play a significant role in the wetland ecosystem. Adult odonates feed on mosquitoes, blackflies and other blood-sucking flies and act as an important biocontrol agent of these harmful insects. In the urban areas of Thailand, larvae of the container breeding dragonfly, Granite ghost (*Bradinopyga geminata*) was successfully used to control *Aedes* mosquito, an important vector of the dengue fever. Many species of odonates inhabiting in agro ecosystems play a crucial role controlling pest populations.

In addition to the direct role of predators in ecosystem, their value as indicators of quality of the biotope is now being increasingly recognised. For example, in South Africa it has been shown how species assemblages of dragonflies change with levels of human disturbance. On the other hand, species recorded at industrial land or urban areas with disturbed riparian vegetation were generalists with wide habitat preference and distribution. These studies also show that dragonflies are sensitive not only to the quality of the wetland but also to the major landscape changes, especially changes in the riparian zone. Recent studies on dragonfly ecology from Western Ghats indicate families like Bamboo tails, Reed tails, Glories, Torrent darts, Torrent Hawks and Club tails are good indicators of health of riverine ecosystem.

Though the Indian odonate fauna is well described in terms of adult taxonomy, their ecology is poorly known. Larval stages of only 76 Indian species are known and the full life history is documented for only 15 species. A good understanding of larval ecology is crucial for odonate conservation. The paucity of ecological information is a serious lacuna when designing any conservation measure. The impact of landscape changes going on since last fifty years or so in the peninsular India on dragonfly distribution and status is not known. This can be tackled only by fresh field surveys to know the threat status and distribution of many species. Future studies on dragonflies may be directed to have a comprehensive understanding of their ecology and their value as a biomonitoring tool. There is no comprehensive account of Indian odonates after Fraser’s fauna volumes published during 1930’s. An assessment by IUCN Red Data Books (International Union for Conservation of Nature, 2004) lists *Burmagomphus sivalikensis*, *Cephalaeschna acutifrons* and *Epiophlebia laidlawi* as threatened Indian odonates. All the three species are restricted to North East India. However a large number of endemic odonates are threatened due to large scale habitat destruction. For example, *Myristica bambootail* the monotypic damselfly of the Western Ghats is restricted to Myristica swamps of evergreen forests. The swamps are very restricted geographically within the ghats. The swamps are being drained in an unprecedented scale for agriculture expansion, especially for the arecanut plantations. Draining of the swamps have caused irreversible damage to the breeding habitat of this species. The case of *Myristica bambootail* is only one example. About 67 species of peninsular Indian odonates are endemic. Most of these species are restricted to the riverine ecosystem. Large scale habitat alterations such as damming channel diversion, sand mining and pollution is seriously threating the survival of these species. Long term conservation of odonates and other freshwater biota can only be assured through appropriate national level policy interventions and definite freshwater biodiversity conservation programmes.

Severe floods in the state of kerala during the monsoons of the year 2018 and 2019 was widely reported. Several studies on ecological impacts and biodiversity of floods in kerala are in progress. In this biodiversity survey of ours, we focused on documenting anisopterans in flood prone areas of tirurangadi taluk. We believe that such a repository of anisoptera diversity may help in assessing the impacts on biodiversity by natural calamities like that of floods.

## Materials and methods

The study was conducted for a period of three months spanning from September to December of 2019. Dragon flies were collected by hand picking method and with the aid of nets. Capturing photos of flies was done on sunny days and during two time periods of the day in which odonates are previously reported to be most active. Photos were captured using Canon 6D camera. For all measurements, we used fine threads and 15 cm-rulers. All graphs were plotted using Microsoft office excel 2007 version. Identification of flies was done with the aid of “Pictorial handbook on common Dragonflies and Damselflies” of K.G.Emiliyamma, M.J.Palot and C.Radhakrishnan and by contacting experts in the field of odonate taxonomy. Number of individuals was estimated by using Line transect method. The method involved the following steps:

I. Set up a string transect line along the study area.
II. Estimate the population abundance by observing either sides of transect line. Shannon- Weiner diversity index was calculated by the standard formula, H = −Σ p_i_ ln p_i_ using Microsoft Excel 2007.Simpson’s diversity index was calculated by using the standard formula

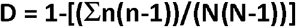

Description of study sites are provided below:

## Description of Study Sites

### Site 1

#### PSMO College

The PSMO College campus (11.05°N and 75.93°E) is situated at Tirurangadi at a distance of 12 km from the Calicut University campus and 24 km from the district headquarters of Malappuram district. It has an average elevation of 10m (33ft.).The vegetation mainly comprises deciduous species, shrubs, herbs and grasses. The area around the campus is enriched with various aquatic habitats like ponds, paddy fields, reservoirs and terrestrial habitats like primary and secondary invasions. Its temperature varies from 28.9°C to 33°C.

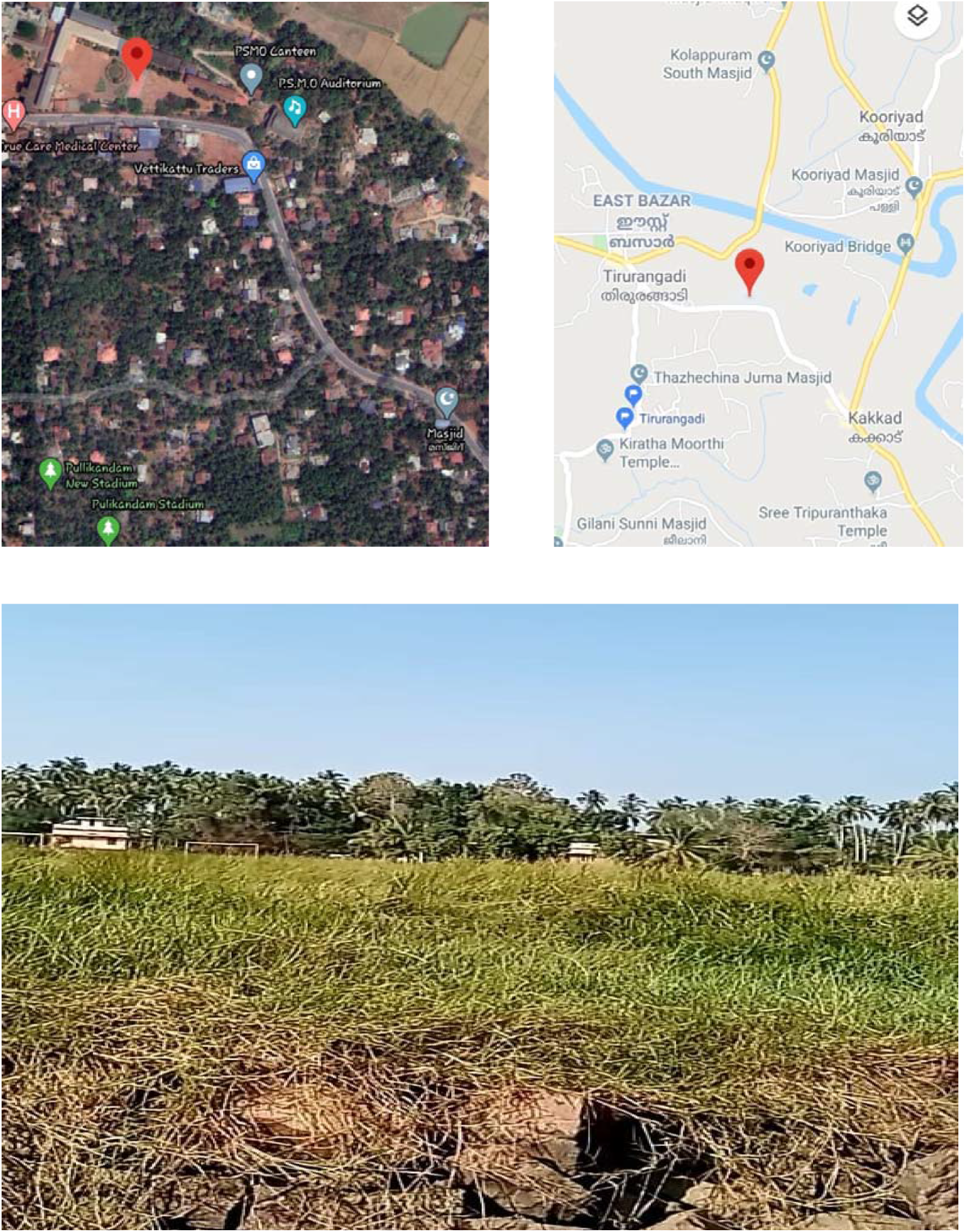

### Site 2

#### Kooriyad

The Kooriyad wetland, (11°2’43”N and 75°56’25”E) is located in Malappuram district along NH 14.A seasonal wetland, Kooriyad is inundated from the onset of monsoon to the end of northeast monsoon. It is a vast area having cultivated, uncultivated and also uninhabited regions that has a diverse flora and fauna. The area around the Kooriyad is enriched with Panampuzha River.

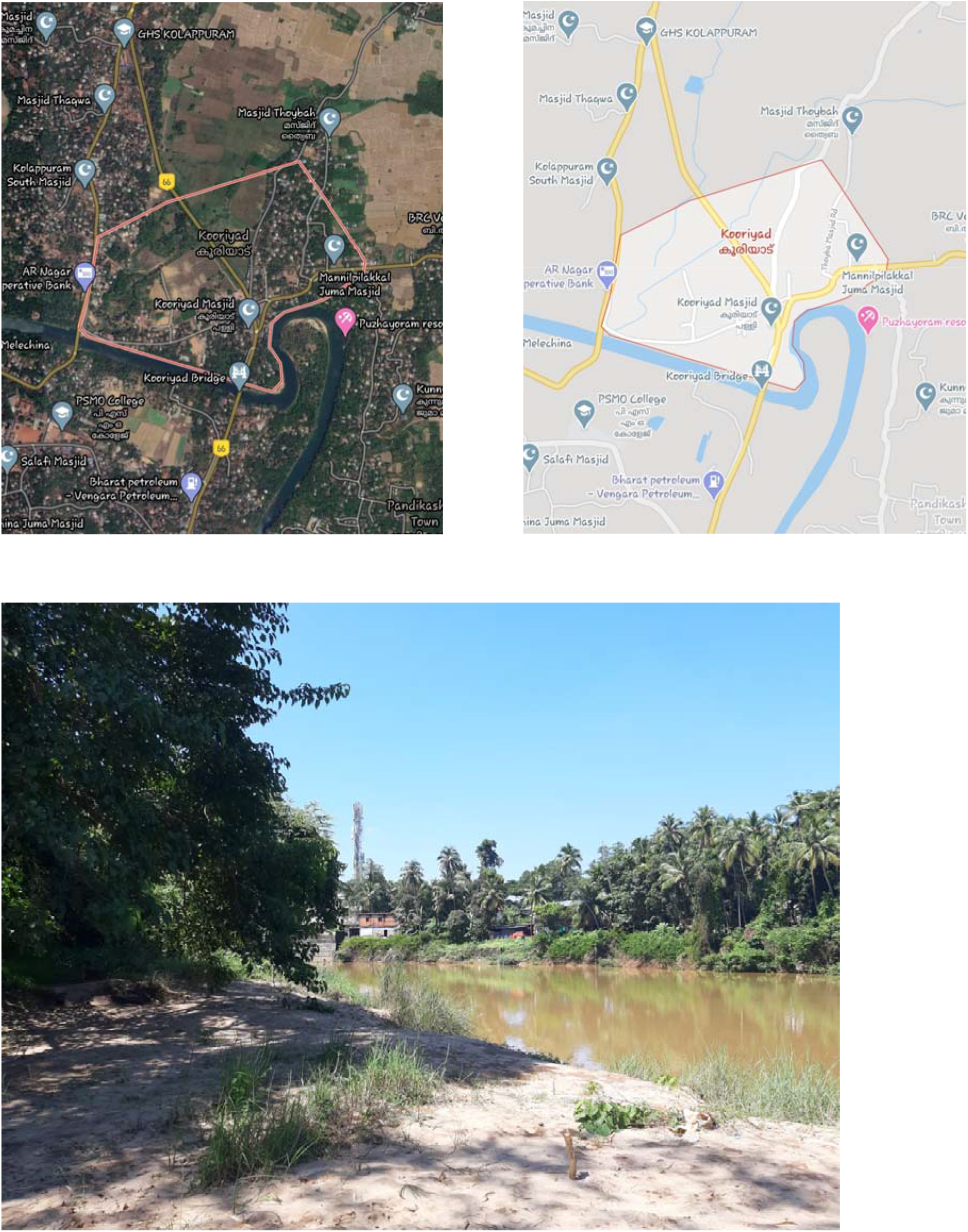

### Site 3

#### Mampuram

Mampuram is located 11°02’40.8”N and 75°55’05.5”E, is a muslim pilgrimage centre located 26 km east of the Tirur.It is situated on the river of Kadalundi. It has a rich diversity of flora due to location on the banks of the river.

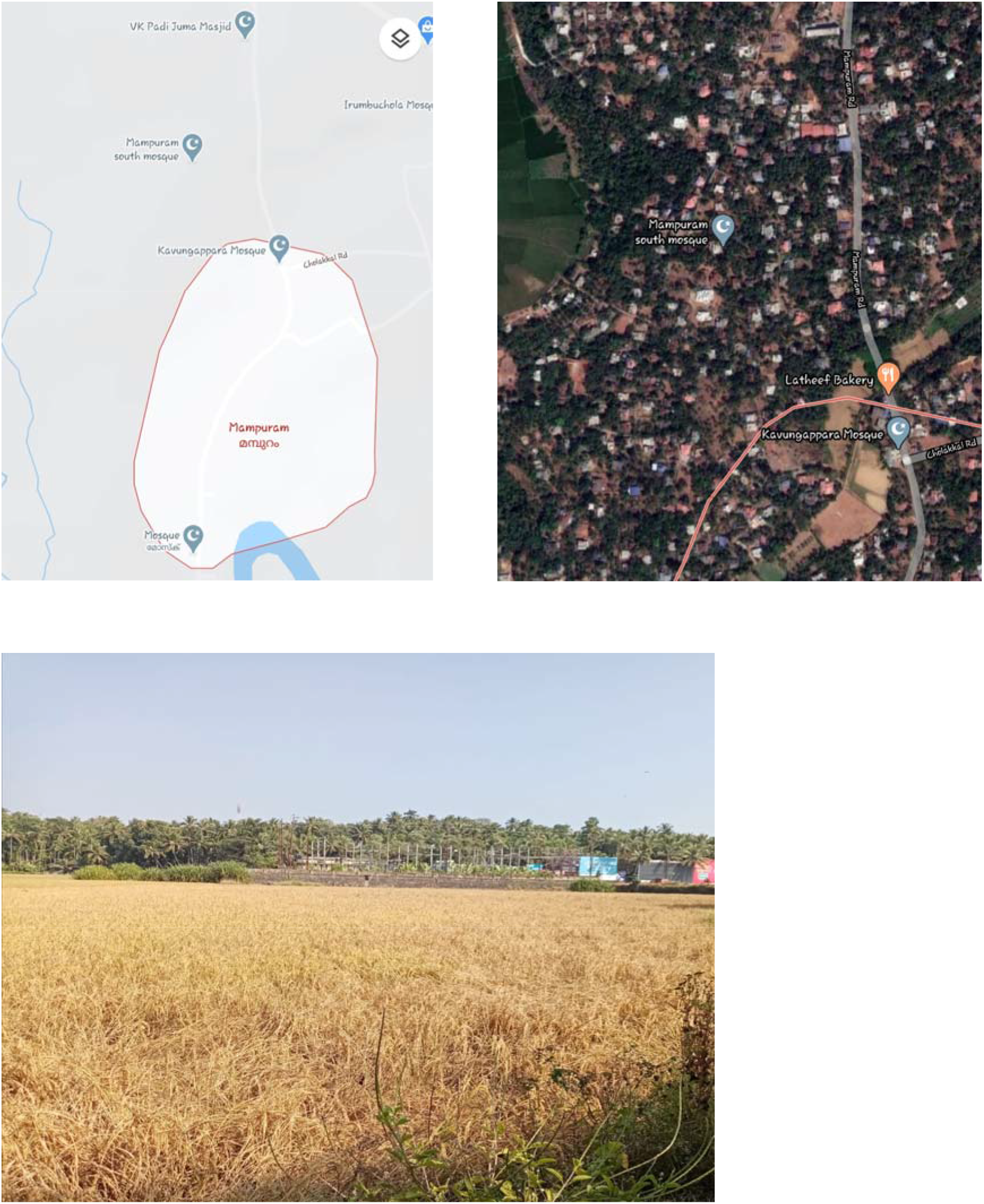

## Results and Discussion

A total of 12 species belonging to 10 genera and 2 families were recorded during the entire study. Sub-order Anisoptera were represented by the family Libellulidae and Aeshnidae, comprising 11 species of Libellulidae and 1 species of Aeshnidae.

Both sub-orders are among the largest dragonflies on the planet. In them, hind wings are broader near the base than the front wings but, forewings are either longer or equal to the length of hind wings. They can fly forwards or backwards or hover like a helicopter. Both are common in fresh water habitats worldwide.

In Libellulidae, many species have pattern wings and distinctive colours on the thorax and abdomen and in Aeshnidae, the abdomens are long and thin. Most are coloured blue and or green, with black and occasionally yellow. Libellulidae can be recognised by a notch on the posterior margin of the eyes and a foot shaped anal loop in the hind wing and Aeshnidae can be recognised by their large, hemispherical, compound eyes which touches the midline and nearly cover their heads. It can be concluded that Libellulidae is representing Odonate family from Kerala (Emiliyamma, 2005). From the collected data “*Neurothemis tullia, Orthetrum sabina, Pantala flavescens* and *Diplacodes trivalis”* were most abundantly found in all the habitats.

By comparing Simpson’s Indices of the study areas, it is evident that Mampuram has highest species richness among the study areas. Among those areas, the PSMO College Campus has the lowest Simpson Index, though all the differences are of small magnitude. Similar trend is also shown for the Shannon Wiener index for the study areas. Higher the Shannon Weiner index, higher is the species richness.

| Sl No: | Scientific Name | Kooriyad | Mampuram | PSMO Campus |
| --- | --- | --- | --- | --- |
| 1 | *Brachydiplax chalybea* | ✕ | ✕ | ✓ |
| 2 | *Brachythemis contaminata* | ✓ | ✕ | ✓ |
| 3 | *Crocothemis erythraea* | ✓ | ✓ | ✓ |
| 4 | *Diplacodes trivalis* | ✕ | ✓ | ✓ |
| 5 | *Neurothemis fulvia* | ✓ | ✕ | ✓ |
| 6 | *Neurothemis tullia* | ✓ | ✓ | ✓ |
| 7 | *Orthetrum sabina* | ✓ | ✓ | ✓ |
| 8 | *Pantala flavescens* | ✓ | ✓ | ✓ |
| 9 | *Rhyothemis variegata* | ✕ | ✓ | ✓ |
| 10 | *Urothemis signata* | ✕ | ✕ | ✓ |
| 11 | *Trithemus festiva* | ✕ | ✓ | ✓ |
| 12 | *Gynacantha dravida* | ✓ | ✕ | ✕ |

## Description of Reported Species

### *Brachythemis contaminata* (Fabricius,1793)

The hindwing of male is 21 to 23 mm in length and the total body length ranges from 29 to 31 mm. The wings of the male are tinted with deep amber except at the tip, and have orange vein and pterostigma. Older males have an orange abdomen. The female is light brown with clear wings and yellowish orange wing spots (pterostigma). The immature male is like the female.

### *Neurothemis fulvia* (Drury,1773)

A medium sized rusty coloured dragonfly with transparent wing tips. In male, abdomen has 21-26 mm and hind wing has 27-32 mm in length. In female, abdomen has 20-24 mm and hind wing has 26-32 mm in length. In male, thorax and abdomen is reddish brown in colour and wings are opaque dark reddish brown with an irregular triangular transparent area at the tip of the wing. In female, colour of head, thorax and abdomen pales than males or rusty brown. Wings are clear amber yellow with a dark ray extending to the tip in fore wings.

### *Neurothemis tullia* (Drury,1773)

It is known as pied paddy skimmer. In male, abdomen has 16-20mm and hind wing has 19-23 mm in length. In female, abdomen has 16-19mm and hind wing has 20-23mm in length. In male, phase is black, thorax is black with mid dorsal cream stripe and abdomen is black with a broad mid dorsal creamy white stripe on the upper side. Wings basal half is opaque blue black which is bordered by a milky white patch towards the tip and its tips are transparent. The female differ significantly from the male in body marking and colouration. Thorax is greenish yellow with a bright yellow mid dorsal stripe. This stripe is broadly bordered with blackish brown throughout. Abdomen is bright yellow with a broad black band above and under side is black. The base of the wings are bright amber yellow and its front edge is blackish brown,broadening into a very large blackish brown spot.This spot extent to the rear edge of the wing. In hind wings, this spot is irregular or sick- le shaped.Tips of all wings are broadly blackish brown.

### *Orthetrum sabina* (Drury,1770)

In male, abdomen is 30-36mm and hind wing is 30-36mm in length. Their phase is yellow- ish green,thorax is greenish yellow with black tiger like stripe and in abdomen the segments 1-3 are green with broad black rings and distinctly swollen at the base. Wings are transparent, inner edge of hind wing tinted with yellow. Female is very similar to the male but abdomen is 32-35mm and hind wing is 31-35 mm in length.

### *Pentala flavescens* (Fabricius,1798)

A medium sized dragonfly with rusty thorax and yellow abdomen. Their abdomen is 29-35 mm and hind wing is 38-40 mm in length. In male, face is bright golden yellow or orange and thorax is olivaceous or rusty and is coated thickly with fine yellowish hair. On sides,it is pale green or blue green,abdomen is bright reddish brown and is tinted with brick red dorsally. The segments 8-10 have black spots above. Wings are bight reddish brown. Female is very similar to male. Eyes are olivaceous brown above and wings are evenly smoky. The abdomen lacks the dorsal red colouring found in the males.

### *Rhyothemis variegata* (Linnaeus,1768)

A medium sized dragon fly with butterfly like yellow and brown wings. In male,abdomen 23-25mm and hind wing is 33-36mm in length. Male are frons iridescent green and abdomen is black. The fore wing is transparent and golden yellow. The wing tip,leading edge and centre of the wing are marked with deep coffee brown spots. The hind wing also has similar spots;however the central spot is absent. Moreover,the wing base is marked with an irregular brown patch. The trailing edge of the hind wing has a characteristic ‘W’ shaped coffee brown mark. In females,the tip of the fore wings is transparent. A dark brown opaque area extends to the centre of fore wing. The area boarders a bright yellow hockey stick shaped patch. In hind wings,the brown opaque area is more extensive and reaches up to the wind tip,which encloses a long yellow central patch and a small yellow spot towards the wing tip. This patch also boarders yellow spots of wing margins and abdomen is bluish black.

### *Diplacodes trivialis* (Rambur,1842)

A small greenish yellow or blue dragon fly with black markings. In male,hind wing is 19-22mm in length. In female,hind wing is 22-24mm. Abdomen is 18-20mm in length. In male,thorax is greenish yellow or oliaceous. The dorso lateral area is violet brown and is speckled with minute dots. In old adults,thorax is covered with fine blue purinescence. The abdomen contains the seg- ments 1-7 greenish yellow with mid dorsal and sub dorsal black stripes. Remaining segment is black and transparent. Female resemble young or sub adult male.

### *Brachydiplax chalybea* (Brauer,1868)

The male of the species is 33 to 35mm long and has a hindwing 24 to 27mm long. It is powder blue with light brown sides and a dark tip to the abdomen. Wings are hyaline, with tinted burnt-brown base, fading to amber. The female is brownish yellow in color with darker markings along the dorsal abdomen. Its wings lack the yellow tinge. This species can be easily distinguished from other species in this genus by its larger size, characteristic colour of the thorax and bases of wings.

### *Crocothemis servilia* (Drury,1770)

It is a medium sized blood-red dragonfly with a thin black line along the mid-dorsal ab- domen. Its eyes are blood-red above, purple laterally. Thorax is bright ferruginous, often blood-red on dorsum. Abdomen is blood-red, with a narrow black mid-dorsal carina. Anal appendages are blood-red. Female is similar to the male; but with olivaceous-brown thorax and abdomen. The black mid-dorsal carina is rather broad.It breeds in ponds, ditches, marshes, open swamps and rice fields.

### *Urothemis signata* (Rambur,1842)

It is a medium-sized dragonfly with red eyes. thorax and abdomen. Its wings are transparent with a amber colored spot surrounded by a dark-brown patch in the base of hind-wings. Its abdomen is blood-red, with some black marking on the dorsum of segments 8 and 9. Female is similar to the male; but yellowish in color. The black spots on the dorsum is repeated on segments 3 to 7. Juvenile and sub-adult males also have these marks.This species breeds in ponds and slow flowing rivers, typically in lowland areas. The males often found perch on exposed twigs. This species has managed to colonise urban water bodies and park ponds.

### *Trithemis festiva* (Rambur, 1842)

Black stream glider is a medium-sized dragonfly with purple color on its body structure. In the male, the frontal area appears darker purplish grey. The eyes are dark brown above, with a purple coloured tinge, which is bluish grey, lateral and beneath. The thorax is black, covered with purple pruinescence, which helps it appear deep blue. The legs are black and wings are transparent with a dark opaque brown mark at the base of hind wing, with a black spot on tip of the wing. The abdomen is covered with fine blue pruinescence. The female looks brown in the front and extends above. The eyes are dark brown above and appear more grey-ish below. Thorax is greenish-yellow to olivaceous, with the presence of a medial dark brown lateral stripe. In addition, Y-shaped inverted stripes can be observed on the sides. Legs are black with anterior femora being yellow on the inner side. Wings are transparent with dark reddish-brown tip with a black spot, similar to the male. The abdomen appears bright yellow with medial, lateral and ventral stripes, coloured black, however, the medial and lateral black stripes form a confluence at abdominal segments to enclose a wedge-shaped yellow spot.

### *Gynacantha dravida* (Lieftinck, 1960)

It is a large dragonfly characterized by its homogeneous colouring of dull browns and greens, by its long and thin anal appendages, and by its crepescular habits. Its principal food ap- pears to be mosquitoes and microlepidoptera. During the day it rests in dark thickets. Fully matured specimens have bright colours; blues and greens developing very late in life. Young specimens have brown color with some dark shades. Females are exactly similar to the males in colors and markings.

It is very closely related to *G.subinteerrupta* and it is difficult to distinguish them. But the relative lengths of the superior and inferior anal appendages are different. The inferior being more than one-third the length of superiors in *G. dravida* and less than one-third in *G. subinterrupta*.

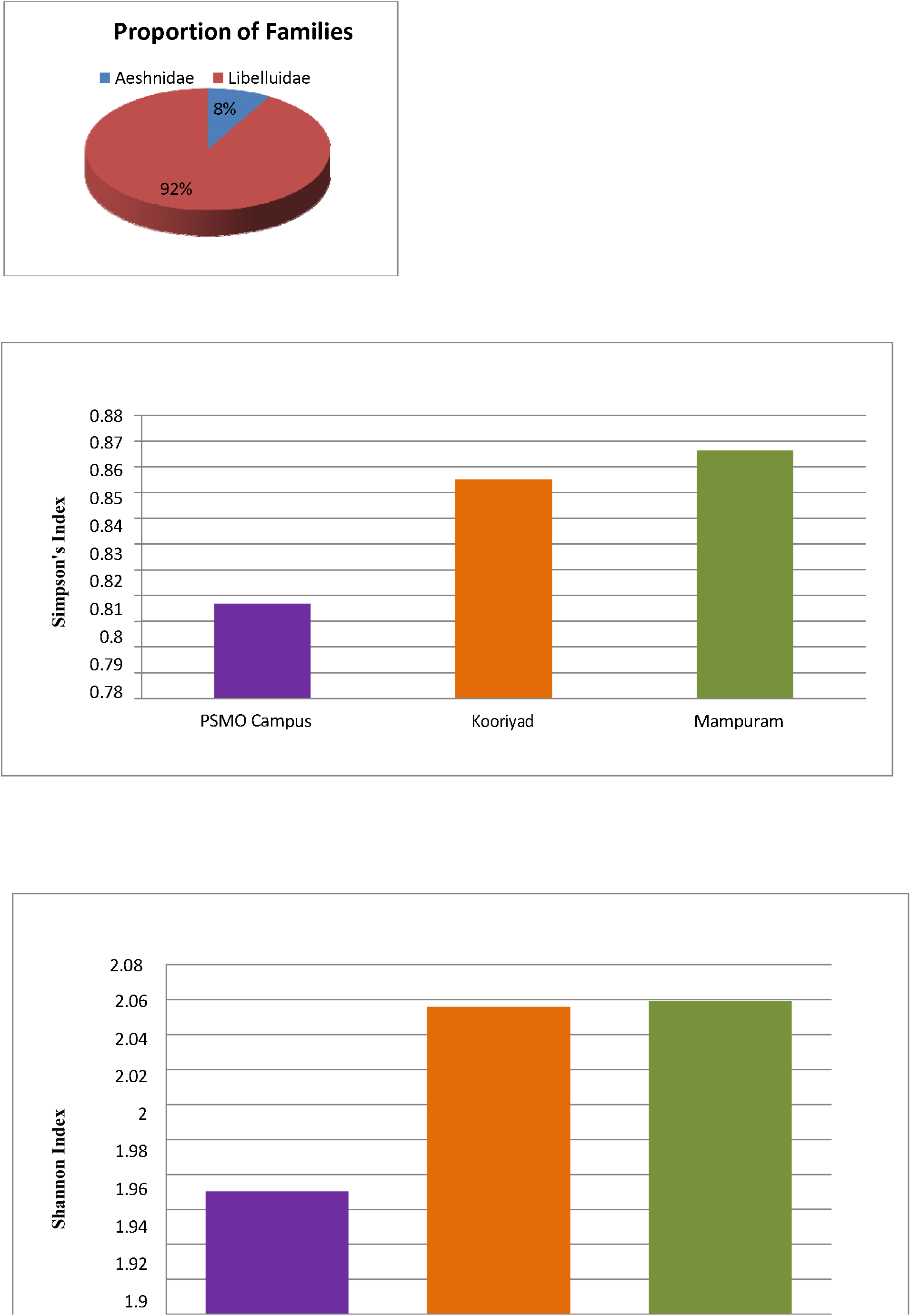

**PLATE 1.**
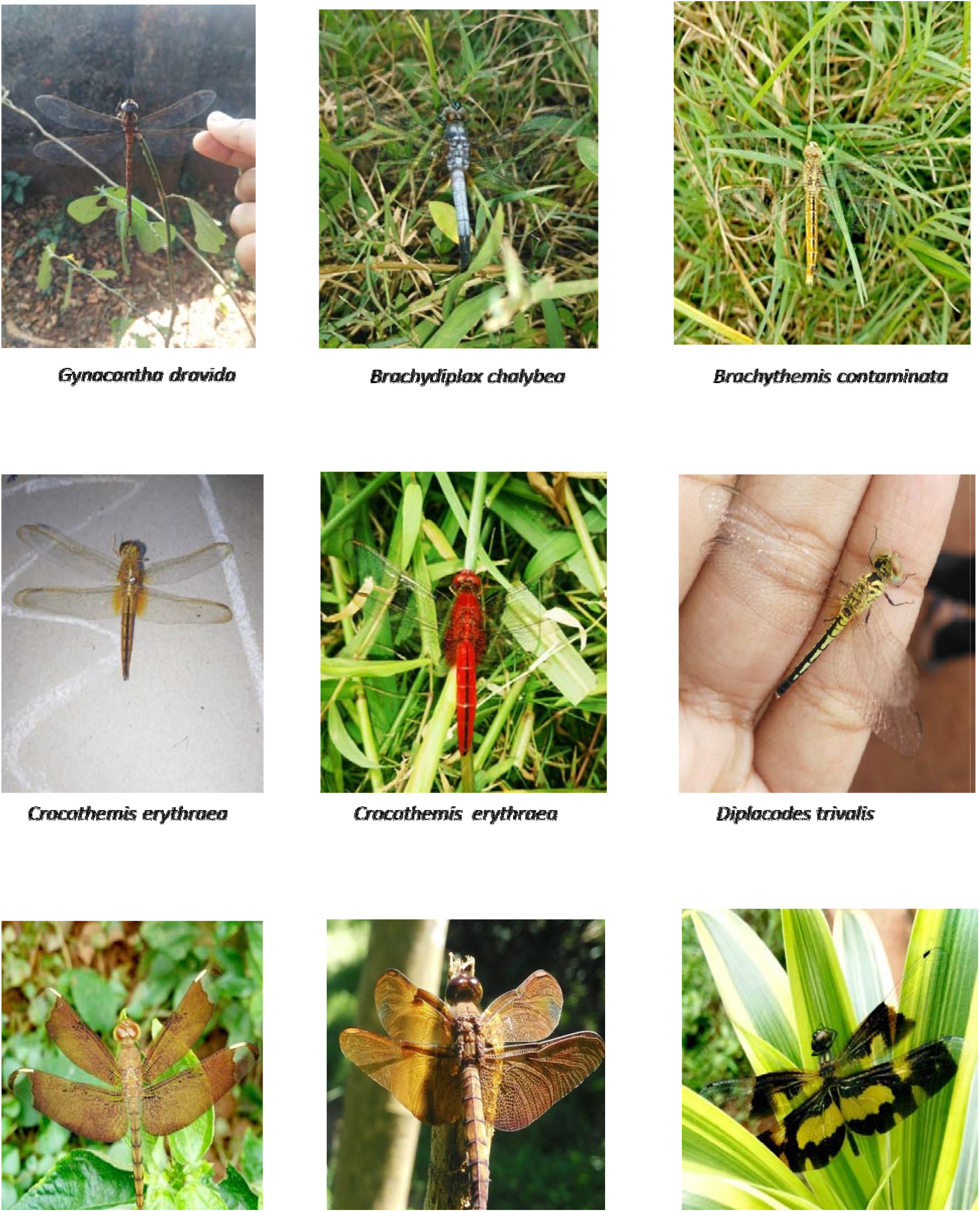

**PLATE 2.**
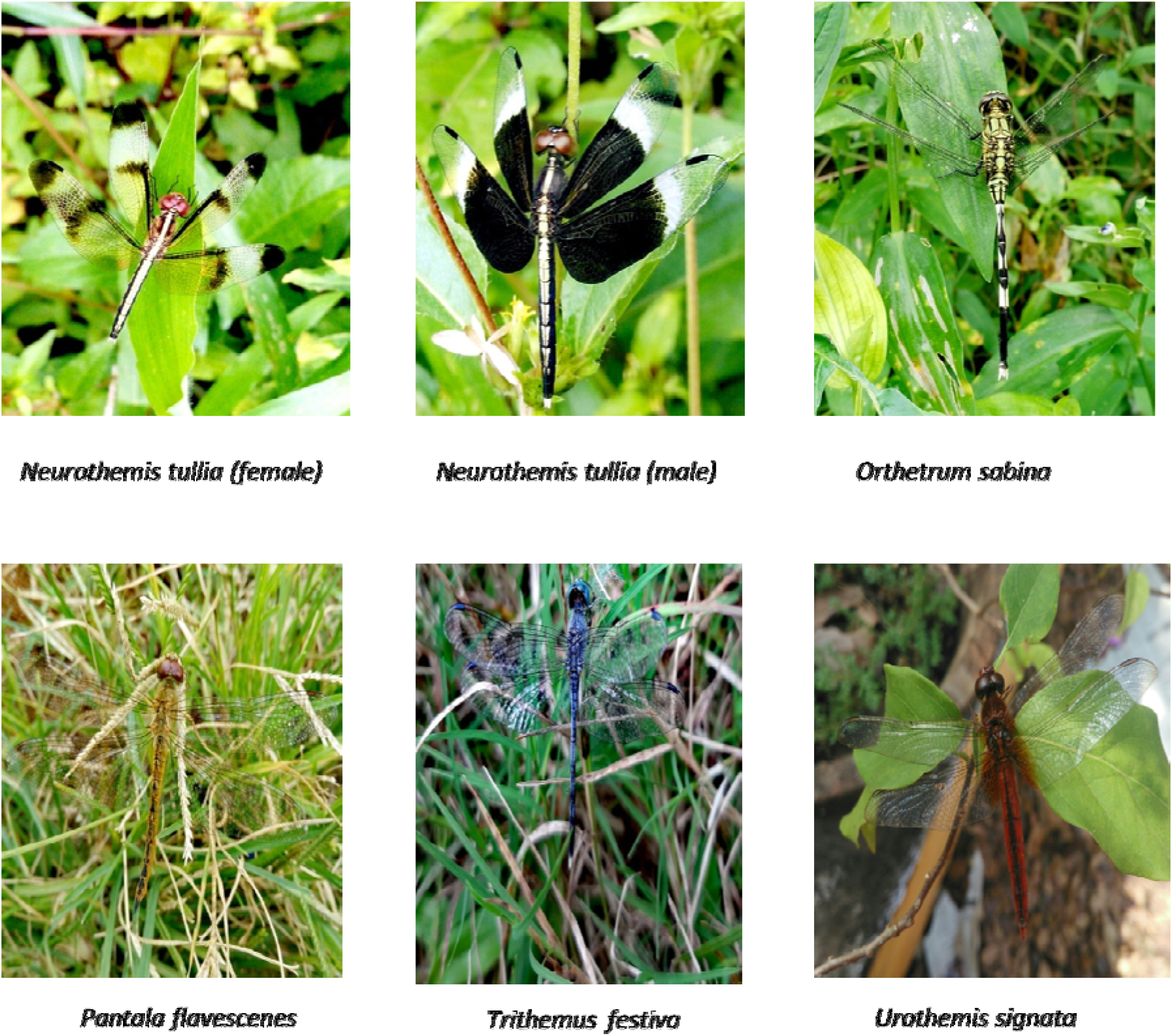

## Notes

### Competing Interest Statement

The authors have declared no competing interest.

